# Visual Motion Detection Thresholds Can Be Reliably Measured During Walking and Standing

**DOI:** 10.1101/2023.06.19.545323

**Authors:** Stephen DiBianca, John Jeka, Hendrik Reimann

## Abstract

In upright standing and walking, the motion of the body relative to the environment is estimated from a combination of visual, vestibular and somatosensory cues. Associations between vestibular or somatosensory impairments and balance problems are well established, but less is known about how the ability of the visual system to detect motion affects balance control. Here we asked whether motion detection thresholds can be reliably measured during walking and standing. Typically, motion threshold values have been obtained while sitting, with the head fixated to eliminate self-motion. In this study we 1) tested whether a visual motion detection threshold can be reliably measured during standing and walking in the presence of natural self-motion; and 2) investigated whether visual motion detection thresholds differ during standing and walking.

**Methods:** Twenty-nine subjects stood on and walked along a self-paced, instrumented treadmill inside a virtual environment displayed on a large dome. Participants performed a 2-alternative forced choice experiment in which they discriminated between a counterclockwise (“left”) and clockwise (“right”) rotation of a visual scene projected on a large dome. A 6-down 1-up adaptive staircase algorithm was implemented to change the amplitude of the rotation. A psychometric fit to the participants’ binary responses provided an estimate for the detection threshold

**Results:** We found strong correlations between the repeated measurements in both the walking (R = 0.84, p < 0.001) and the standing condition (R = 0.73, p < 0.001) as well as good agreement between the repeated measures with Bland-Altman plots. Average thresholds during walking (mean = 1.04 degrees, SD = 0.43 degrees) were significantly higher than during standing (mean = 0.73 degrees, SD = 0.47 degrees).

**Conclusion:** Visual motion detection thresholds can be reliably measured during both walking and standing, and thresholds are higher during walking.

## Introduction

Vision plays an important role in balance control for standing and walking by providing information about movement relative to the environment via optical flow [1]. Quantifying the capabilities of the human visual system is challenging, as any particular property such as contrast sensitivity [2], depth perception [3,4], and motion detection [5] may affect different functional behaviors. The ability to detect self-motion from optical flow is expected to be most relevant for balance control, but motion detection is typically measured during sitting [6–10], where self-motion is eliminated, and balance is not an issue. Our motivation for this study was to investigate whether visual motion thresholds can be reliably measured during standing and walking, and whether they might be relevant for balance control in different behavioral contexts.

When studies investigate the relationship between visual processing and fall risk, they typically use measures of visual acuity such as contrast sensitivity [11,12], depth perception [13,14], or size of the visual field [15,16]. Visual acuity is meaningful for maneuvering around an environment and avoiding falls caused by tripping or hitting obstacles [15] as vision provides information about object size, location, and where to place the swing leg foot into a safe space. Visual acuity relates to central vision, or focal vision, capable of high spatial resolution and particularly useful for pattern and object recognition [17]. While visual acuity mostly concerns central vision, visual motion perception is more related to peripheral vision [6]. Illusion of self-motion in response to visual motion has been shown to be primarily influenced by stimuli in the peripheral visual field [18]. Optic flow can produce illusions of self-motion, and thus disturb upright balance in both standing [19–21] and walking [22–25]. To our knowledge, there has been only one study that has directly compared measures of visual acuity to motion perception in their relationship to control of upright balance. Data collected during the Salisbury Eye Evaluation (SEE Project) [8] found that in a model including visual acuity, contrast sensitivity, visual field, and motion detection threshold, the motion detection thresholds were associated with over 3 times higher odds of failing on a single leg balance stance task when adjusted for age, sex, and race compared to the other measures of vision. A review by Saftari and Kwon [26] highlights the general finding that decreased visual acuity is associated with increased risk for falls and hip fractures. Despite these findings, they emphasize that visual motion perception as a contributor to fall risk has been a critical omission in the literature.

Here we tested the reliability of the participants’ visual system in detecting optic flow through a visual motion detection test using psychophysical methods. Visual motion detection tests have typically been performed while sitting [6–10]. Here we measured visual motion detection thresholds during standing and walking, where body sway generates a natural background level of self-motion. A threshold measure characterizes the underlying mechanism [27] of a sensory system. In the case of a visual motion detection test, this threshold provides a measure of how sensitive the visual system is in detecting movement in the environment. To calculate a threshold value, perceptual responses are recorded after exposing participants to optic flow with varying directions and amplitudes. In this study, we use a common adaptive psychophysical method, in which the amplitude of the stimulus is increased or decreased depending on the history of responses. Our hypotheses are that 1) visual motion detection thresholds are correlated between repeated measures in both standing and walking and 2) thresholds in walking are different than in standing.

### Research Design and Methods

Twenty-nine healthy participants (14 female, 39 ± 15 years old) between the ages of 23-67 were recruited for this experiment. Subjects provided informed verbal and written consent to participate. Subjects did not have a history of neurological disorders or surgical procedures involving the legs, spine, or head within 6 months of the protocol. The experiment was approved by the University of Delaware Institutional Review Board.

### Experimental Protocol & Setup

Participants stood and walked on a self-paced, tied-belt treadmill (Bertec, Columbus, OH, USA) surrounded by a virtual environment displayed on a large dome that occupied the subjects’ full visual field (Figure 1). All participants started with a 15-minute walking block to familiarize themselves with the environment, walking on the self-paced treadmill, and the two-alternative forced choice task (2AFC). After the familiarization block, participants performed four blocks of the 2AFC task, alternating between standing and walking, with the order counter-balanced between participants, with 15 participants walking first and 14 standing first. In the standing blocks, participants stood 2 meters way from the center of the curved screen. Six reflective markers were placed on the subjects: two on the temples, two on the occipital condyles, and two on the posterior superior iliac spines. Marker positions were recorded using a Qualisys Motion Tracker System with 13 cameras at a sampling rate of 200Hz. For self-paced control of the treadmill, a nonlinear proportional-derivative (PD) controller was implemented via Labview (National instruments Inc., Austin, TX, USA) to keep the midpoint of the two markers on the posterior iliac spine at the midline of the treadmill. Visual perspective and position in the virtual world were linked to the midpoint between the two markers placed on the subjects’ temples and superimposed over the forward motion dictated by the treadmill speed. Subjects wore a safety harness in the event of a fall, although none occurred.

**Figure 1:**
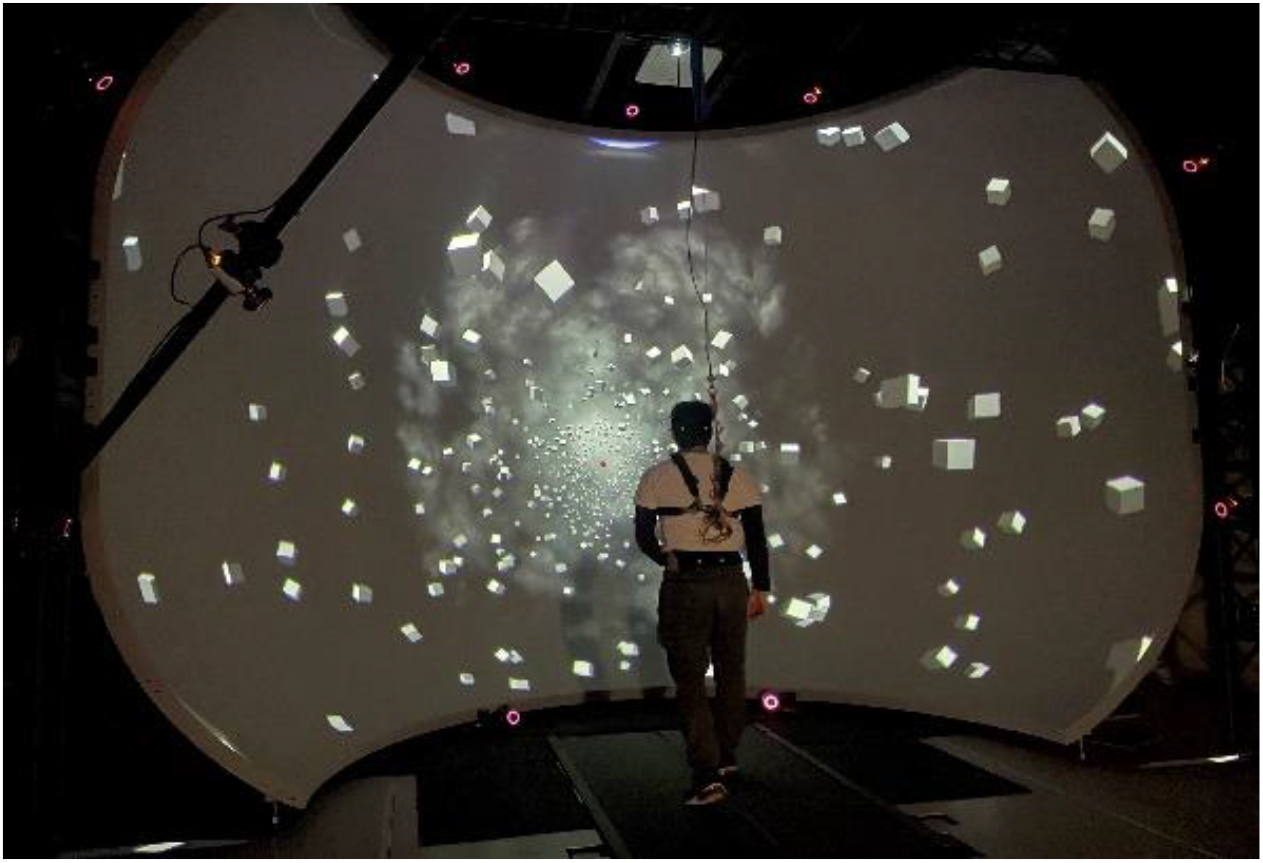
Experimental setup depicting a participant walking in front of the virtual reality dome on the self-paced treadmill.

### The Virtual Scene

The experimental setup is displayed in Figure 1, showing a participant walking on the self-paced treadmill in the virtual environment designed and implemented in Unity3d (Unity Technologies, San Francisco, CA, USA.). The scene consisted of 1,000 cubes floating before a dark background, randomly distributed in a cylindrical tunnel along the anterior-posterior axis with a radius of 14-40 m from the central axis through the treadmill. Each cube was 1×1×1 m in size. A red sphere was linked to the midpoint between the two markers placed on the temples as a focal point for participants and was placed 50 m ahead. Fog was displayed in the distance to obfuscate the end of the tunnel and create the perception of infinite distance. The anterior-posterior movement of the virtual scene matched the speed of the treadmill.

### Two-alternative forced choice task

The 2AFC task presents participants with a rotation of the virtual environment around the anterior-posterior axis of the treadmill in a counter-clockwise (left) or clockwise motion (right). Participants verbally reported the direction of motion as “left” or “right”. The stimulus waveform was a single cycle of a raised cosine for velocity with a frequency of 1 Hz. Each block consisted of 100 trials, where one trial is a single stimulus. After each response, the experimenter initiated the next trial. After every 25 trials the subject was given a brief break to release concentration on the task, then indicated when ready to continue, which typically took about 15 seconds. In the walking trials, subjects kept walking normally during these breaks. After each block of 100 trials, subjects took longer breaks of at least two minutes, more if needed.

### Adaptive Staircase for stimulus amplitude

The amplitude of the stimulus was the angle of rotation around the anterior-posterior axis. The amplitude was adjusted using an adaptive 6 down 1 up staircase algorithm [28] for parameter estimation by sequential testing [29]. The amplitude decreased after six correct responses and increased after one incorrect response, until 100 trials were completed. Figure 2 shows an example of the adaptive staircase from one single block consisting of 100 trials. The initial amplitude was 4 degrees.

**Figure 2:**
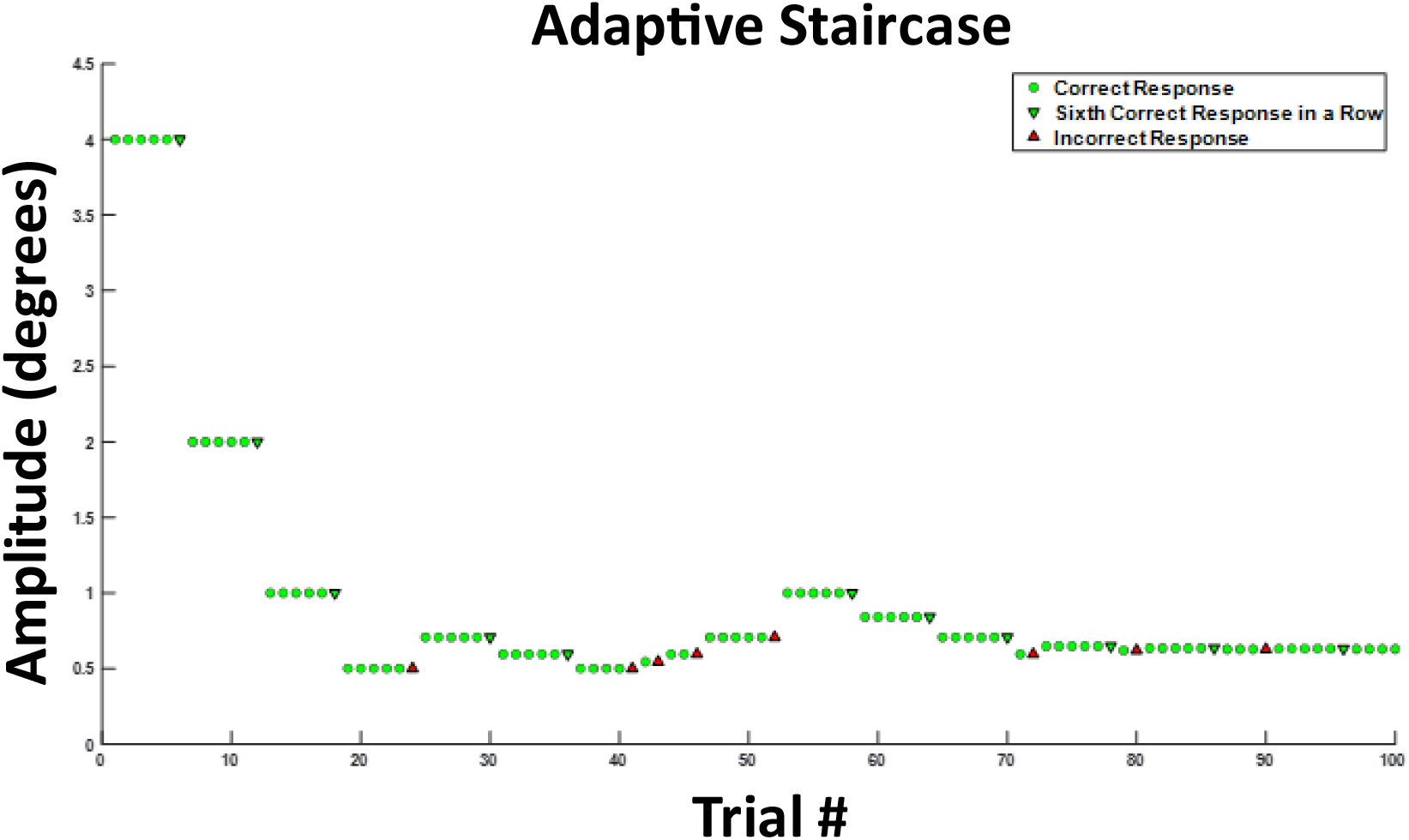
Exemplary data from a participant performing the 2-alternative forced choice task. The green circles represent correct answers after which the stimulus amplitude stayed the same. A green downward arrow represents a 6^th^ correct response in a row, which leads to a decrease in stimulus amplitude, making the task more difficult. An upward red arrow represents an incorrect response, which leads to an increase in movement amplitude, making the task easier. This is a 6-down-1-up adaptive staircase.

### Psychometric Fit

To obtain a visual motion detection threshold, we fit a psychometric curve to the 100 binary responses of the 2-alternative forced choice task in each condition. The fit was performed in MATLAB’s fitglm function using a generalized linear model (GLM) with a probit link. The motion detection threshold was defined as the value corresponding to a target probability of 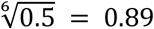 [28–31]. Figure 3 displays a psychometric fit for one participant during one of the conditions. The threshold value is highlighted by the dotted red line that corresponds to the amplitude of rotation at which the subject responded “right” with 89% confidence.

**Figure 3:**
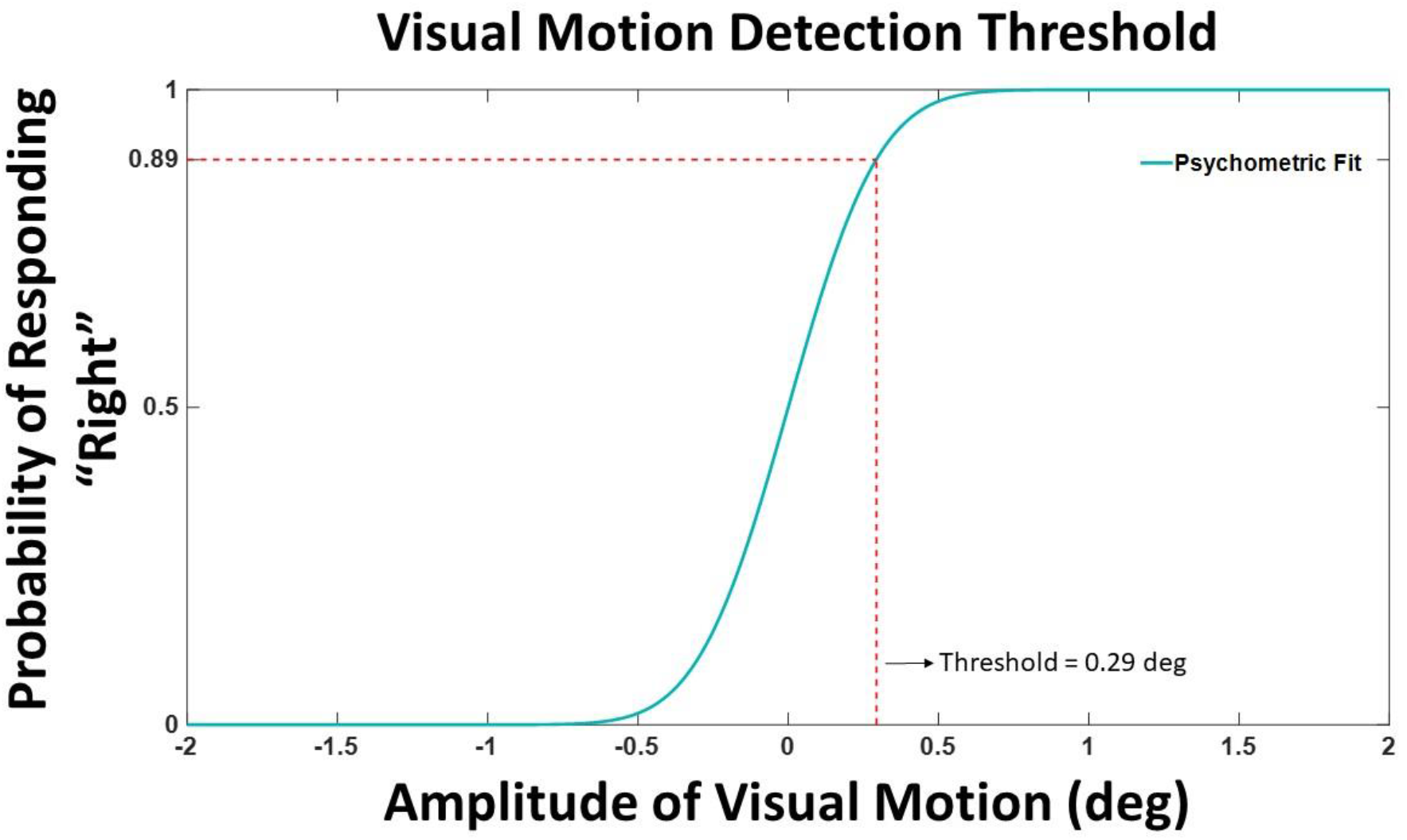
Example psychometric fit data from one participant (VMD04). The threshold value is defined by the angle in degrees at which the participant was responding with a rightward rotation with 89% confidence. The amplitude of the rotation is displayed on the horizontal axis, with positive values corresponding to rightward rotations and negative values to leftward rotations. The probability of responding “right” is on the vertical axis.

The mean and standard deviation of the underlying normal distribution represent the bias and the slope for the psychometric fit. A bias value of 0 indicates an equal chance of left and right guesses (50%) at 0 amplitude movement. The bias of the psychometric curve was set to 0 for all participants. The slope of the psychometric fit is determined by the standard deviation of the underlying gaussian distribution, which characterizes the acuteness of detection, or how accurate the visual system can detect the stimulus, visual motion [32]. Exemplary fits for one participant (VMD04) are shown in Figure 5. Here the slope of the walking trials (dark blue) is steeper than the slope of the standing trials (cyan), resulting in a larger threshold or a less accurate ability to detect motion in the environment while walking.

### Statistical Analysis

The normality and homoscedasticity of the visual motion detection thresholds for both walking and standing per block and per condition were evaluated using a Shapiro-Wilk test of normality and an F-test. While the walking threshold data met these assumptions, the standing threshold data did not. Parametric tests were used for the walking data while any comparisons made with standing included non-parametric tests.

For Hypothesis 1 on the agreement between threshold measurements obtained in trial one and two, Pearson’s correlation coefficients were calculated between the two measurements of both walking and standing conditions. We used Bland-Altman plots to show the mean difference between the repeated measures and construct limits of agreement [33]. A paired, two-tailed t-test was performed to test for significant differences between the repeated measures for walking block one and two. A paired samples Wilcoxon test was performed to test for significant differences between the repeated measures for standing block one and two. To test Hypothesis 2 whether a difference exists between visual motion thresholds during walking and standing, a paired samples Wilcoxon test was performed between the thresholds measured during walking versus standing, averaging across the two repeated measures for each subject.

## Results

All subjects completed the experiment of both walking and standing conditions. On the individual level, 24 subjects had higher thresholds during walking versus standing. The average walking speed for participants during the walking condition was 1.09 m/s (SD = 0.20 m/s).

### Test-Retest Reliability

Thresholds were obtained from walking block one (mean = 1.13 degrees, SD = 0.45 degrees), walking two (mean = 0.97 degrees, SD = 0.41 degrees), standing one (mean = 0.73 degrees, SD = 0.41 degrees) and standing two (mean = 0.74 degrees, SD = 0.54 degrees). Both walking and standing conditions showed a strong positive correlation between measurements one and two. The correlation coefficient between the walking measurements was 0.84 (p < 0.001), and the correlation coefficient between the standing measurements was 0.73 (p < 0.001). Figure 4 displays threshold values from trial one and two against each other for both walking (Panel A) and standing (Panel B). Also shown in Figure 4 are the Bland-Altman plots for walking (Panel C) and standing (Panel D). The mean absolute difference between measurement one and two for walking was 0.16 degrees and the mean difference between standing measurements one and two was 0.01 degrees. Results of the t-test between walking blocks showed a significant difference between the repeated measures (p = 0.001). Results of the Wilcoxon test between standing blocks did not show a significant difference between the repeated measures (p = 0.59).

**Figure 4:**
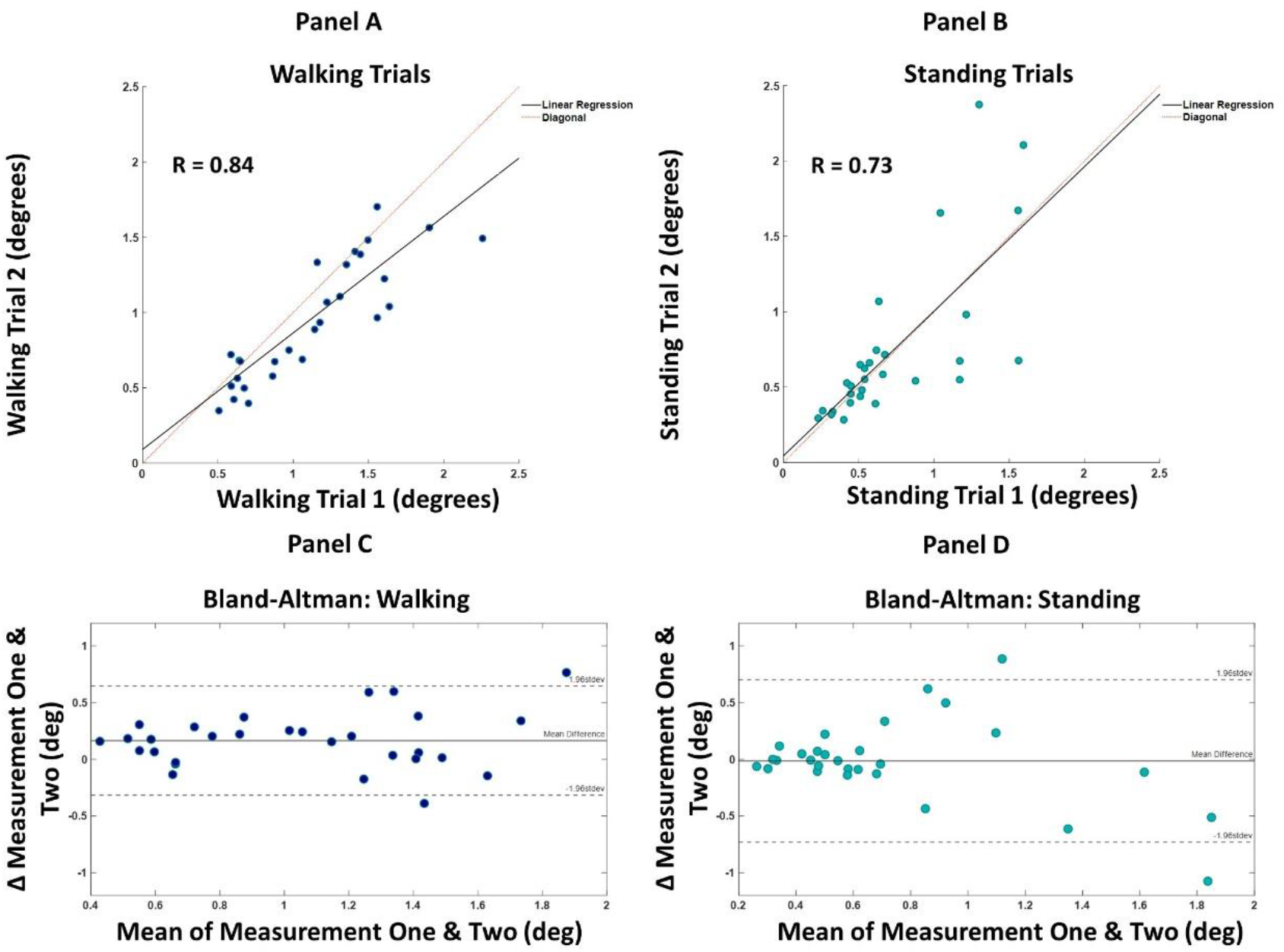
Panel A shows the visual motion threshold values for walking, with block one on the horizontal and block two on the vertical axis. The dotted red line is the diagonal, and the black line is the linear regression. Panel B shows the same for standing. Panel C shows the Bland-Altman plot for walking, with the mean of the two blocks on the horizontal and the difference on the vertical axis. Solid horizontal lines indicate the mean difference and dashed horizontal lines mark two standard deviations from the mean. Panel D shows the same for standing.

### Walking Versus Standing Visual Motion Thresholds

We observed higher visual motion detection thresholds in walking vs. standing. Figure 5 (Panel A) shows the visual motion detection thresholds for walking (dark blue) and standing (cyan). Thresholds were significantly higher during walking compared to standing (p < .001), with an average threshold value of 1.04 degrees (SD = 0.43) for walking and 0.73 degrees (SD = 0.47) for standing. (Figure 5, Panel A). Panel (B) shows psychometric fits from an exemplary participant for trials one and two for walking and standing. The slopes in the two standing blocks are steeper than in the two walking blocks.

**Figure 5:**
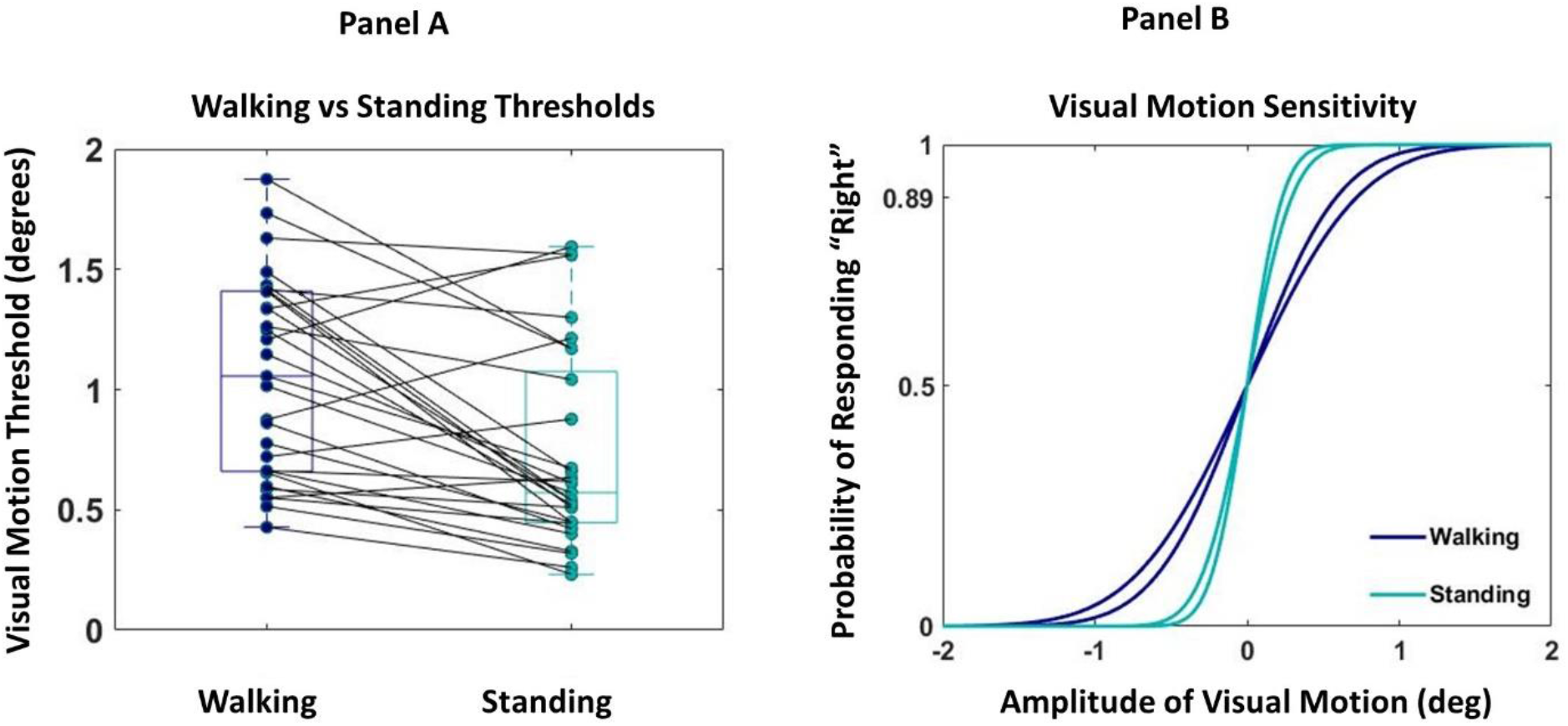
Panel (A) shows box and whisker plots of visual motion thresholds for walking versus standing. Boxes and whiskers are the median, quartiles and inter-quartile ranges. Dots are data from individual participants, where each dot is the average between the two repeated measures. Gray lines connect walking and standing measures from the same participant. Panel (B) shows exemplary psychometric fits for participant VMD04 for both walking and standing. The horizontal axis represents the stimulus amplitude, with positive sign corresponding to rightward rotation, and the vertical axis represents the probability that the participant responded with “right”.

## Discussion

Our study investigated the reliability of measuring a visual motion detection threshold during walking and standing and compared these measures between those two tasks. We used virtual reality to display a visual scene that rotated in a clockwise or counter-clockwise motion at different amplitudes and asked participants to report the direction of rotation. We found that a visual motion threshold can be reliably obtained during tasks involving body sway and movement, namely standing and walking. We also found evidence that visual motion detection thresholds are higher when walking versus standing, potentially influencing the way visual information is processed between the two tasks.

Visual motion detection thresholds can be reliably obtained during walking and standing. Strong correlations indicate good agreement between the two measures taken at different time points. Bland-Altman plots in Figure 4 support this finding by indicating small average differences between the two measurements for both walking (0.16 degrees) and standing (0.01 degrees). However, it should be noted that a significant difference did exist between the two walking trials, where thresholds in walking block one were systematically higher than in walking block two. This difference could be the result of an adaptation to the experiment. Some subjects noted that they started to change strategies towards the end of the experiment by focusing less hard on the red dot and paying attention to the periphery to detect the motion, without losing fixation of the red dot. This systematic difference was not observed between standing blocks (p = 0.59). Visual motion thresholds while walking are on average higher than in standing, indicating larger visual movement is required to detect changes in the visual surround during walking than in standing.

Traditionally, visual motion detection tests are performed while sitting, with the head fixated to avoid any type of head movement [6–10]. Our motivation was to perform the psychometric test in an ergonomic manner that would quantify visual motion thresholds during natural movements of the head and body while maintaining upright balance, opposed to a restrained position. Since participants were not restrained in any way, potential cues from the vestibular and proprioceptive system from the natural motion may have influenced the participants’ threshold results. Although small, a visual stimulus can provoke a response that may cause movement of the head and/or ankles, cueing the vestibular or proprioceptive system. Some subjects noted that they reported perception of self-motion when the visual stimulus became small, rather than movement of the visual stimulus. Such cues from other sensory systems may have influenced responses that could not be considered purely visual. Head movements in standing are relatively small, but walking produces considerable head movement and might influence a person’s ability to detect motion depending on the direction in which the stimulus is moving relative to the head. For the walking conditions, the stimulus was manually triggered by the experimenter with a 1-2 s time window between stimuli regardless of the phase of the gait cycle. For example, a stimulus may have rotated to the right while the participant was moving towards the left or right, resulting in natural and virtual movements either adding up or canceling each other out. The added head movement during walking in conjunction with inconsistencies of stimulus onset during the gait cycle may be a contributing factor to the increased motion detection thresholds observed during walking.

We estimated the detection threshold by fitting the slope of the psychometric curve, i.e. the standard deviation of the underlying normal distribution, to the response data for each participant. Bias, or the mean of the underlying normal distribution, can also be used as a parameter for fitting. A bias can occur if a participant either habitually chose a particular side when they were unsure of the direction or if the visual system itself had a skewed mapping (i.e. left movement more noticeable than right). Including the bias can lead to improper fits in rare cases [34], and it has been found that participants could voluntarily shift their central bias, causing changes in threshold values, but the slope parameter remained relatively unchanged [32]. Since we had no a priori reason to expect a bias, we constrained the mean of the psychometric curve to zero for all participants. To investigate possible effects of this choice, we also repeated our analysis with fitting both slope and bias. We found that with this choice, the correlation between the slope estimates decreased for both the walking trials (R = 0.57, p=0.001) and standing trials (R = 0.67, p<0.001). Additionally, we asked whether the added bias parameter provides meaningful information about the motion perception system for each participant, or rather represents a superfluous parameter that leads to overfitting. To this end, we analyzed the repeatability of the bias estimates between the two trials, using the same approach as for the threshold estimate in the main analysis. We found weak correlation for walking (R = 0.35, p=0.064) and standing (R = 0.38, p=0.041). Although there was significance for the standing condition, there is little consistency between the bias measures of the two repetitions in both walking and standing, indicating that adding bias as a parameter for the psychometric curve fit represents overfitting rather than meaningful information about the visual system.

It may be beneficial to understand when during the gait cycle we are most susceptible to visual influences on balance and thus when a visual motion detection threshold may be most useful to quantify. For example, when a person is in double stance, they receive more information from lower limb proprioceptors and thus may rely less on vision, while moments of single stance, in which the contralateral leg is in swing phase, may utilize visual cues more to maintain balance since there is less contact with the body to the ground. In fact, phase-dependent visual coupling has been observed during walking [22]. One major mechanism for balance control is modulation of the foot placement based on the state of the body at mid-stance [35], and visual motion detection is likely used to estimate the body state. The reliability of motion detection may change during different phases of the gait cycle, although this question needs further investigation and is not addressed in this study. Overall, our results indicate that our ability to detect motion during standing and walking differ regardless of head and body position during stimulus onset.

